# The structure of the enigmatic ripple phase in saturated bilayers resolved: Machine learning reveals four lipid populations

**DOI:** 10.1101/2021.11.25.470048

**Authors:** Matthew Davies, A. D. Reyes-Figueroa, Andrey A. Gurtovenko, Daniel Frankel, Mikko Karttunen

## Abstract

A new mixed radial-angular, three-particle correlation function method in combination with unsupervised machine learning (ML) was applied to examine the emergence of the ripple phase in dipalmitoyphosphatidylcholine (DPPC) lipid bilayers using data from atomistic molecular dynamics (MD) simulations of system sizes ranging from 128 to 4,096 lipids. Based on the acyl tail conformations, the analysis revealed the presence of four distinct conformational populations of lipids in the ripple phases of the DPPC lipid bilayers. The expected gel- (ordered; *L*_o_) and fluid-like (disordered; *L*_d_) lipids are found along with their splayed tail equivalents (*L*_o,s_ and *L*_d,s_). These lipids differ based on their gauche distribution and tail packing. The disordered (*L*_d_) and disordered splayed (*L*_d,s_) lipids spatially cluster in the ripple in the groove side, that is, in an asymmetric manner across the bilayer leaflets. The ripple phase does not contain large numbers of *L*_d_ lipids, instead they only exist on the interface of the groove side of the undulation. The bulk of the groove side is a complex coexistence of *L*_o_, *L*_o,s_ and *L*_d,s_ lipids. The convex side of the undulation contains predominantly *L*_o_ lipids. Thus, the structure of the ripple phase is neither a simple coexistence of ordered and disordered lipids nor a coexistence of ordered interdigitating gel-like (*L*_o_) and ordered splayed (*L*_o,s_) lipids, but instead a coexistence of an ordered phase and a complex mixed phase. Principal component analysis (PCA) further confirmed the existence of the four lipid groups.

## Introduction

The existence of the asymmetric ripple phase was first discovered by Tradieau, Luzzatti and Reman almost 50 years ago.^1^ Later, in 1980, Chen, Sturtevant and Gaffney identified a new phase transition when the ripple phase disappears and the bilayer becomes uniform.^2^ Much of the details of the ripple phase (often called *P*_*β′*_) remained unknown but in a landmark study Sun et al.^3^ analyzed electron density from x-scattering and observed that the saw-tooth shaped ripple has two sides, a major side, often called the major arm, that is gel-like and a fluid-like minor arm. In a related study, Sengupta, Ragunathan and Katsaras argued that the minor arm cannot be in the fluid, or the so-called *L*_*α*_, phase as otherwise the length of the arm would be longer due to the significantly larger area per lipid in the *L*_*α*_ phase.^4^

The structure of the ripple phase has remained debated ever since and the most accurate picture of the ripple is perhaps from the x-ray scattering study of Akabori and Nagle.^5^ As their main results, they reported that the lipids are not in registry in the curved area of the ripple and in the minor arm of the ripple. Furthermore, they hypothesized that in addition to the gel-like ordered (*L*_o_) lipids in the major arm, the minor arm has five different conformational classes of lipids. Lipids are, however, typically classified as ordered and disordered.^6^ As an additional complication, the main phase transition, despite being of first order, has long been known to have features of critical behaviour.^7–9^

Computer simulations and theory have contributed significantly to this debate as well. The first observation of the ripple phase in atomistic MD simulations was reported in 2005 by de Vries et al.^10^ While the asymmetry was present in their MD simulations, in contrast to Sun et al.,^3^ de Vries et al. did not find the co-existence of a gel and fluid but instead they wrote “The organization of the lipids in one domain of the ripple is found to be that of a splayed gel; in the other domain the lipids are gel-like and fully interdigitated”. ^10^ A similar conclusion was reached by Lenz and Schmid using a coarse-grained model. ^11^ More recent atomistic MD studies have reported two-phase co-existence of liquid- and gel-like lipids as well.^6,12^ The molecular level structure and the conformational components that contribute to the ripple remain unresolved.

In this study, we performed atomistic MD simulations using systems of 128, 512 and 4,096 DPPC lipids over 16 *µ*s over a broad temperature range (details in Methods). We then applied conformational clustering based on a three-particle intramolecular density distribution and unsupervised machine learning methods from image analysis to identify conformational lipid classes. The analysis showed the existence of four distinct lipid conformations, ordered, disordered, ordered splayed and disordered splayed. The existence of these classes was then directly verified by analyzing the clusters using principal component analysis (PCA) as well as other methods and they were used to resolve the structure of the ripple phase. The analysis of the ripple shows that it has a complex asymmetric structure in which the two splayed lipid classes have an important role.

## Results

### Three-particle correlation function to measure local structure

To quantify local structure, we used the mixed, radial-angular three-particle distribution function, *g*_3_ ≡ *g*_3_(*r*_*BC*_, *θ*_*ABC*_), originally introduced by Sukhomlinov and Müser in 2020 in a different context.^13,14^ In this method, illustrated in Figure 1, a triplet of atoms is selected: 1) the central atom (blue), *B*, 2) the nearest neighbour to the central atom (orange), *A*, and 3) the other atom within the cutoff (red), *C*. Physically, *g*_3_ is proportional to the probability of finding the other atom at a distance of *r* from the central atom when the vectors 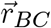 and 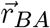 form an angle of *θ*_*ABC*_. Figure 1 gives three examples of this using the same central and nearest neighbour atoms in each case.

**Figure (1).**
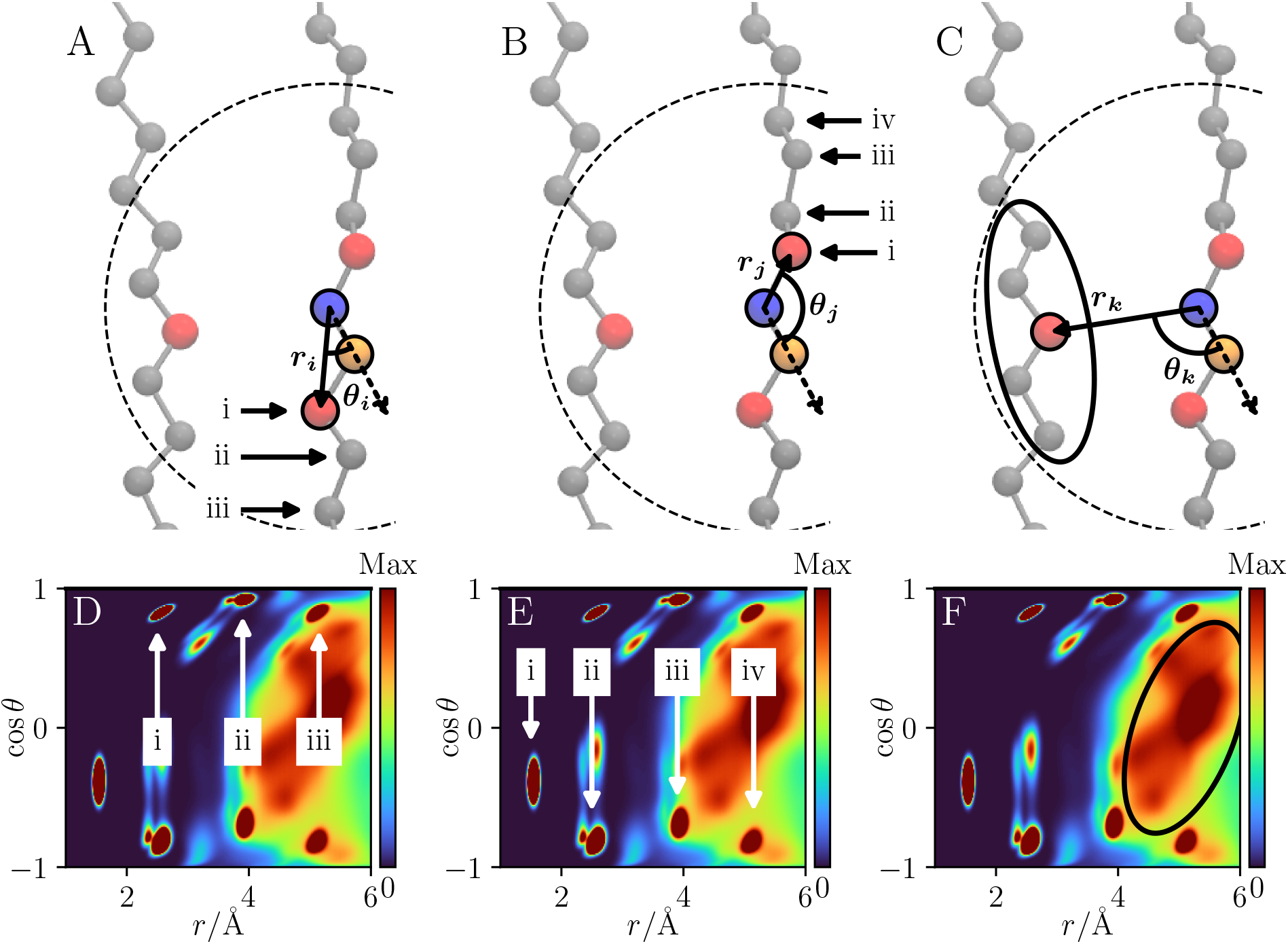
Schematic of the mixed angular-radial three-particle correlation function applied to a single carbon in a DPPC acyl chain/lipid tail. A central atom (blue) and its nearest neighbour atom (orange) are selected. The angle formed between the three atoms with the central atom at the apex, *θ*, and the distances between the central atom and the selected atoms, *r*, are calculated. This is repeated for all target atoms within the cutoff (dashed line). Three key structural contributions are shown here with an example target atom in red. The ‘same-side’ tail (A), ‘opposite-side’ tail (B) and ‘other’ tail (C). Below each diagram the locations of the main contribution to the probability distribution by each of these different tail contributions have been indicated. The same tail contributions are marked with arrows, labelled showing the peak density and contributing atom (from the central atom shown). The ‘other’ tail is much more diffuse and contributions from single atoms are less resolvable so this has been indicated by the black ellipse showing the contributed region. The same-side contributions are radially offset relative to the opposite-side because the nearest neighbour atom is not included.

Here, *g*_3_ was applied on the carbon atoms of the DPPC molecules in two ways: 1) to calculate the bilayer average including both inter- and intra-molecular contributions and 2) to analyze each DPPC molecule individually, only assessing the intra-molecular contributions. For the calculation of *g*_3_, 401 radial and 201 angular bins were used with a cutoff of 7 Å. The cutoff was chosen to be greater than the packing distance^15^ to capture the salient structural details. Above a critical value capturing the nearby chain contributions the analysis was not sensitive to the choice of cutoff value. The nearest neighbour distance was determined using the traditional radial distribution function but other criteria can also be used.

Figure 1 shows three structural details specific to the DPPC carbon atoms. Panels A and B show two cases of bonding within the chosen tail. This is relative to the nearest neighbour; since the nearest neighbour is not included in the distribution, the same-side contributions are shifted. The distinct peaks in the distributions correspond to the atom *n* bonds away from the central atom on the opposite side and *n* + 1 bonds away on the same side. The first peak in both cases picks up the adjacent atom angular component and hence does not split. For *n* ≥ 2, in a bonded chain, there are increased degrees of freedom which allow peak splitting. Panel C shows the packing of other tails. This is the contribution by the carbon atoms not in the same lipid tail as the target central atom, but in another lipid tail packed close to the lipid tail in question.

### Lipid clustering and lipid similarity

To compare the *g*_3_ distributions of individual lipids with each other, the mean structural similarity index metric (SSIM)^16^ was applied. SSIM was originally developed as a metric for determining the similarity of images. Here the images are the *g*_3_ distribution matrices as computed above for each of the individual lipids. SSIM is unique compared to many other metrics in that it combines three quantities to determine the similarity metric and, second, these quantities are evaluated locally.

The three quantities in the original paper were luminance, contrast and structure. ^16^ In the case of lipid *g*_3_ probability distributions, the differences in mean probability, standard deviation, and in spatial probability distribution are used. The structural similarity index metric (SSIM), can be defined in each of the local windows as

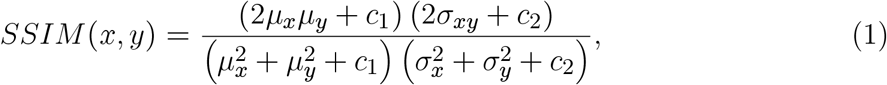

where *x* and *y* refer to the window in each distribution matrix, *µ* is the window mean, *σ*^2^ is the variance of the window, *σ*_*xy*_ is the covariance of the two windows and *c*_1_ and *c*_2_ are small correction factors to avoid division by zero. The final SSIM is the average over the local windows,

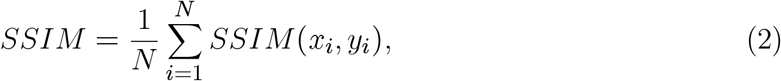

where *x*_*i*_ and *y*_*i*_ are the *i*^th^ window in matrices *x* and *y*, and *N* is the total number of windows. This was applied to all simulated systems. The data range was set to the maximal density value in all compared *g*_3_ distributions. A window size of 7 bins was used. It was verified that the result was independent of the window size.

All lipids (*N*_lipid_) from all temperatures were used to sample the conformational phase space. This generated *N*_lipid_ features (similarity scores) for each of the lipids in the system. The features were combined into an *N*_lipid_ × *N*_lipid_ similarity matrix. The difference here compared to other works is that other ML techniques build feature matrices like this by using predefined distances between predefined atoms.^6,17,18^ Here, the natural length scales emerge spontaneously from the system and its properties via *g*_3_.

### Unsupervised machine learning

After generating the similarity scores, dimensionality reduction was performed by applying the t-distributed stochastic neighbor embedding (t-SNE)^19^ which was scaled to have unit variance. This reduced the similarity matrix to 2-dimensional embedded representation. The unsupervised machine learning clustering algorithm called density-based spatial clustering of applications with noise (DBScan) ^20^ was applied to the 2d data. DBscan was chosen instead of the commonly used *k*-means^21^ clustering since in DBScan the number of clusters is not pre-defined but emerges from the data. Second, and very importantly, in contrast to *k*-means, DBScan is able to find clusters that are non-linearly separable. DBScan needs a neighborhood defined by the parameter *ϵ*. A value of 0.085 was used here but this will vary based on the system analysed. The selection of a suitable *ϵ* is key to identifying all the clusters. This value was selected and verified by using the “elbow plots”,^20^ by visual inspection of the scaled dimensionally reduced data coloured according to the clusters, a plot of the similarity matrix reduced to 2d via principal component analysis (PCA), and confirmed via examination of the *g*_3_ distributions of the lipids. Feature extraction from the clusters was completed by visualizing the *g*_3_ distributions within each cluster to determine the locations of density differences. The lipids within each of the clusters were then identified and further analysed as will be discussed below. Figure 2 summarizes the data analysis and clustering process.

**Figure (2).**
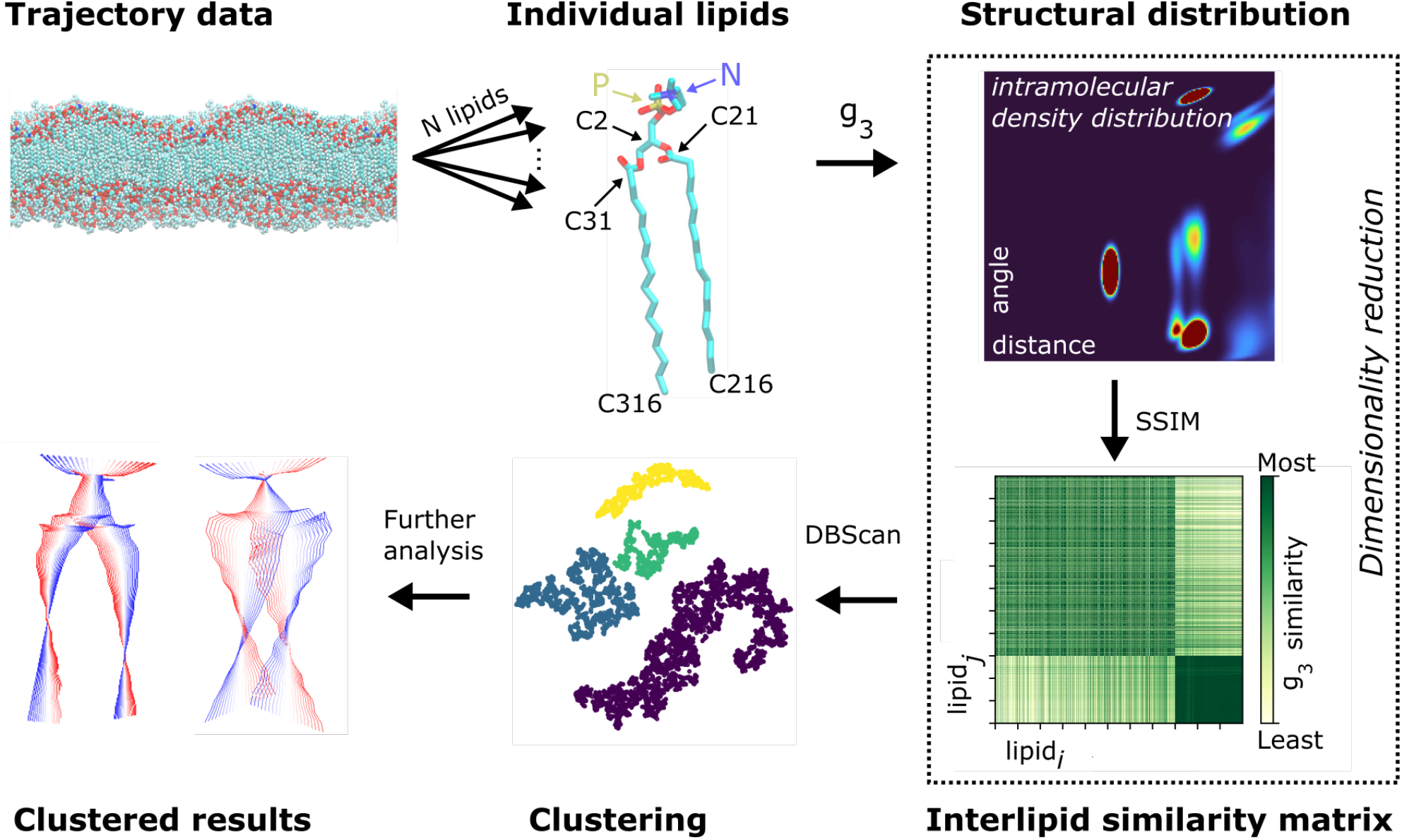
Conformational clustering: First each lipid (some of the important atoms are marked) is isolated from the trajectory and *g*_3_ is calculated to quantify the *intramolecular* structure. The mean structural similarity index metric (SSIM), Equation 2, of each lipid *g*_3_ distribution is compared with all other lipids. Embedding with t-SNE^19^ reduces the matrix of similarity values to a 2d form (from *N*_lipids_ dimensions). This 2d data is clustered using DBScan^20^ to find similar conformational groupings. This is mapped back onto the bilayer for further analysis. The similarity matrix shows how similar the distribution of row *i* is with the distribution of row *j*. The distinct difference in the final quarter of similarity matrix is a result of passing over the phase transition and the large conformation difference this causes.

### Structural details from *g*_3_ analysis and the main phase transition

The particular strength of *g*_3_ is that it is able to detect and quantify structural motifs and changes within the DPPC lipid tails without any a priori knowledge of the underlying lipids, structural metrics or connectivity. The algorithm sees merely the system as a monatomic cloud of atoms. Details such as tail bonding, packing and conformational properties emerge spontaneously in the distributions; this analysis could be applied to any molecule or system undergoing structural changes and help aid in the determination of system structure without prior knowledge of the structural properties relevant to the molecule.

Figure 3 shows temperature dependence of the conventional deuterium order parameter (*S*_CD_) and *g*_3_ using both intra- and inter-molecular contributions. *S*_CD_ measures the ensemble average orientation of the C-H bond vector (*θ*) with the bilayer normal 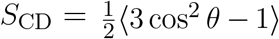. The phase transition temperature detected by these two independent metrics is identical, between 317 K and 318 K. This is slightly above the experimental values of about 314 K^22–24^ but consistent with other simulations using the CHARMM36 force field.^25^

**Figure (3).**
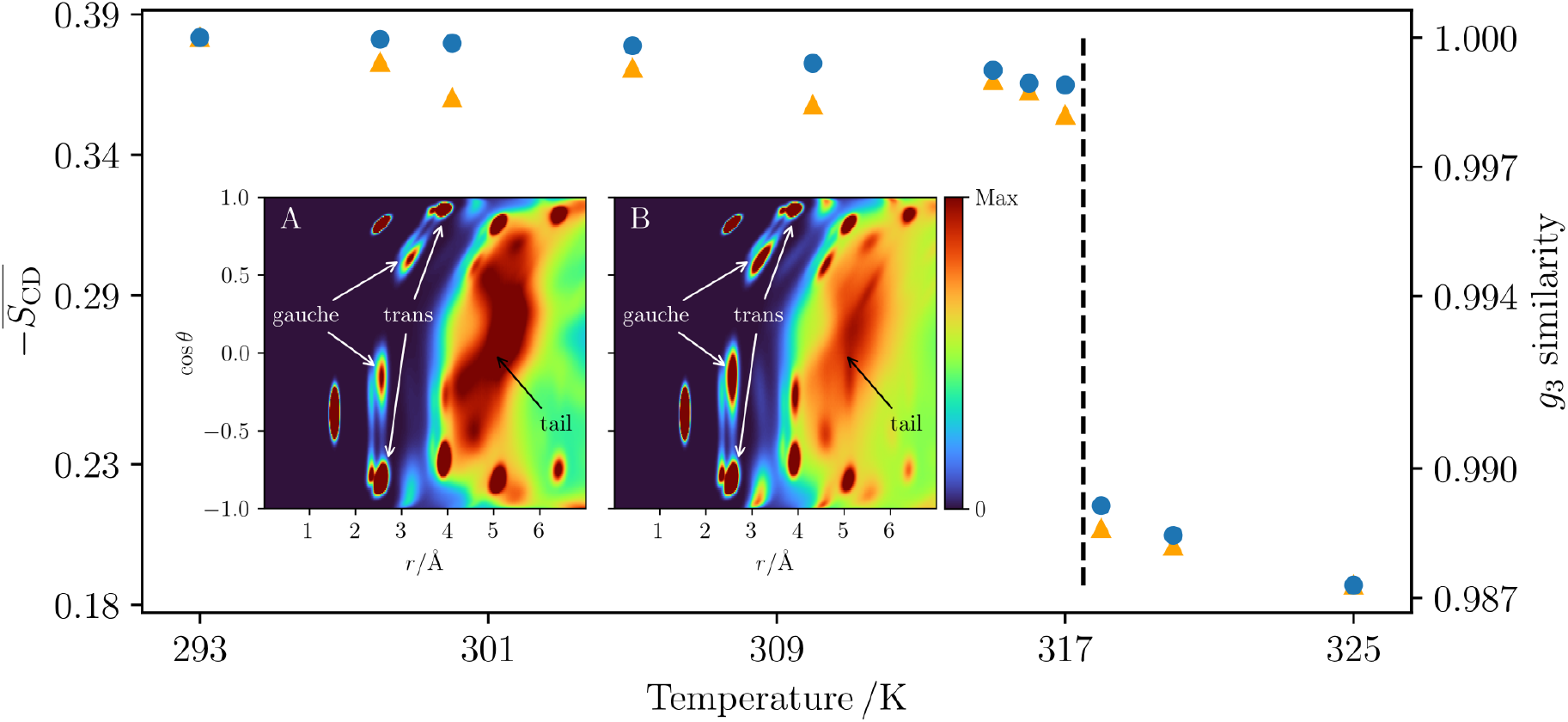
Phase transition detected by the average deuterium order parameter, 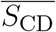, (orange) and using *g*_3_ similarity (blue). Both deuterium order parameter and *g*_3_ similarity (SSIM) are able to detect a phase change between 317 K and 318 K. Insets: Probability density distributions for *g*_3_ of the DPPC carbon atoms at 317 K (A) and 318 K (B). The dihedral gauche and trans peaks have been marked along with the contribution by the other tails.

The inset in Figure 3 shows *g*_3_ distributions below (317 K) and above (318 K) of the phase transition. The distributions display two key features: Firstly, and most obviously, the influence of the other tails on the distribution (see the discussion in connection with Figure 1). Below the transition, the lipid tails are packing in rigid, fixed positions, producing a sharper peak which drops off at larger distances. Above the transition, this peak becomes much more diffuse, with lower peak height and spread across a longer distance, reducing the level of drop off compared to the more ordered lipids.

The second noticeable difference is in the trans/gauche peaks. Below the transition, the gauche peak density is significantly reduced when compared to above *T*_m_. The ordered lipids show less gauche dihedrals in the tails, a well-reported metric for determining the main phase transition. The phase transition can easily be verified even by visual inspection: As the temperature decreases below *T*_m_, it switches from a flat, planar bilayer to a kinked one with regions of different thickness and of asymmetric lengths, Figure 4. This was robust and all bilayer sizes and system temperatures below *T*_m_ adopted a ripple conformation with the exception of one of the 128 lipids systems, which has been noted as a finite size effect.^26^

**Figure (4).**
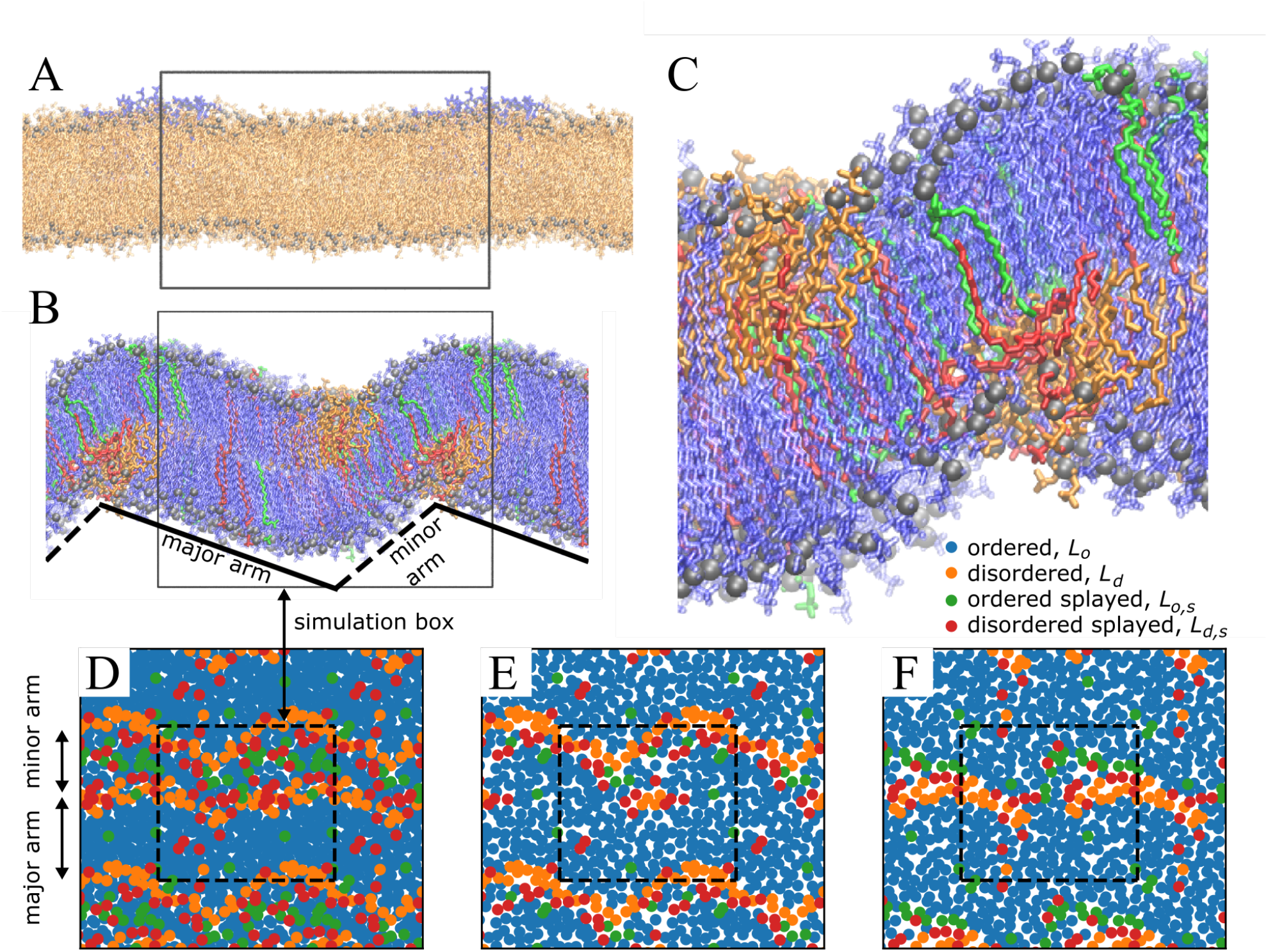
Snapshots from simulations at 318 K (A) and 317 K (B), either side of *T*_m_ (see Figure 3). A zoomed in snapshot of the ripple in the 317 K system (C). All snapshots are coloured according the the conformational cluster the lipid has been assigned. The ripple phase features two regions, the major arm (solid line) and the minor (dashed line). (D)-(F): Top views of the 317 K system. (D) Shows the sum of the two leaflets, (E) and (F) the upper and lower leaflet, respectively. The simulation box is indicated in the figures by the box and periodic images are included for clarity. The two arms are asymmetric in length in agreement with experiments and theory.^1,3,27,28^

### The structure of the ripple phase

While the system-wide *g*_3_ is capable of detecting the phase transition on a system-wide scale, the ripple phase is not homogeneous. The structure of the ripple phase can, however, be analyzed by examining the lipid-wise *g*_3_ and comparing the interlipid similarity. Importantly, clustering (using DBScan) revealed *four* distinct conformational motifs for lipids: disordered (*L*_d_), ordered (*L*_o_), ordered-splayed (*L*_o,s_) and disordered-splayed (*L*_d,s_). These structural motifs emerged in both the 512 and the large 4,096 lipid systems but only three of them appeared in the 128 lipid system, with no differentiation between the ordered-splayed and disordered-splayed lipids; as recently pointed out by Walter et al., finite size effects can be significant in lipid systems.^26^

An example of each of these lipid conformations and the *g*_3_ distribution of this is shown in Figure 5. The unsplayed-splayed behaviour is captured by the lipid tail contribution, and the ordered-disordered behaviour by the gauche defects.

**Figure (5).**
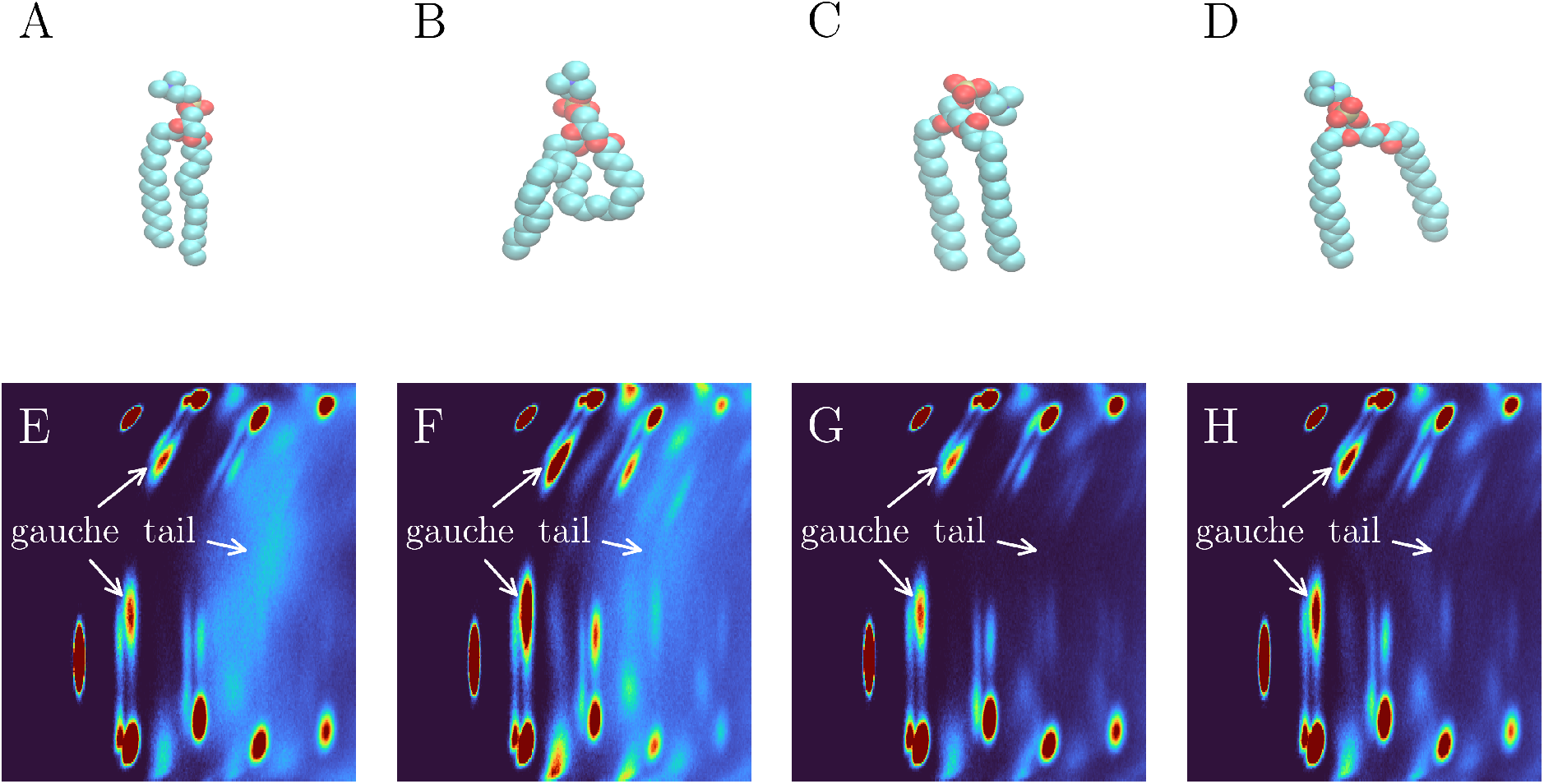
Exemplary lipid from each of the identified clusters taken from the 315 K simulation. A linear-ordered lipid (A), disordered lipid (B), splayed-ordered lipids (C) and splayed-disordered lipids (D). The corresponding *g*_3_ distributions are also shown. E, F G, and H correspond to A, B, C, and D respectively. The first gauche defect peaks (up and down tail) have been marked with arrows and the contribution of the other tail peak has been marked with an arrow for clarity.

The conformational dynamics of each of the lipids clusters was assessed with lipid PCA.^29^ Figure 6 shows the main component of each of the lipid clusters. The disordered lipids’ (*L*_d_ and *L*_d,s_) main motion is a scissoring motion, which has been seen in the fluid phase DPPC lipids in a range of force fields.^29^ The ordered (*L*_o_ and *L*_o,s_) lipids instead have their tails constrained and primarily have a twisting rotational motion. All of the four components are fully distinct.

**Figure (6).**
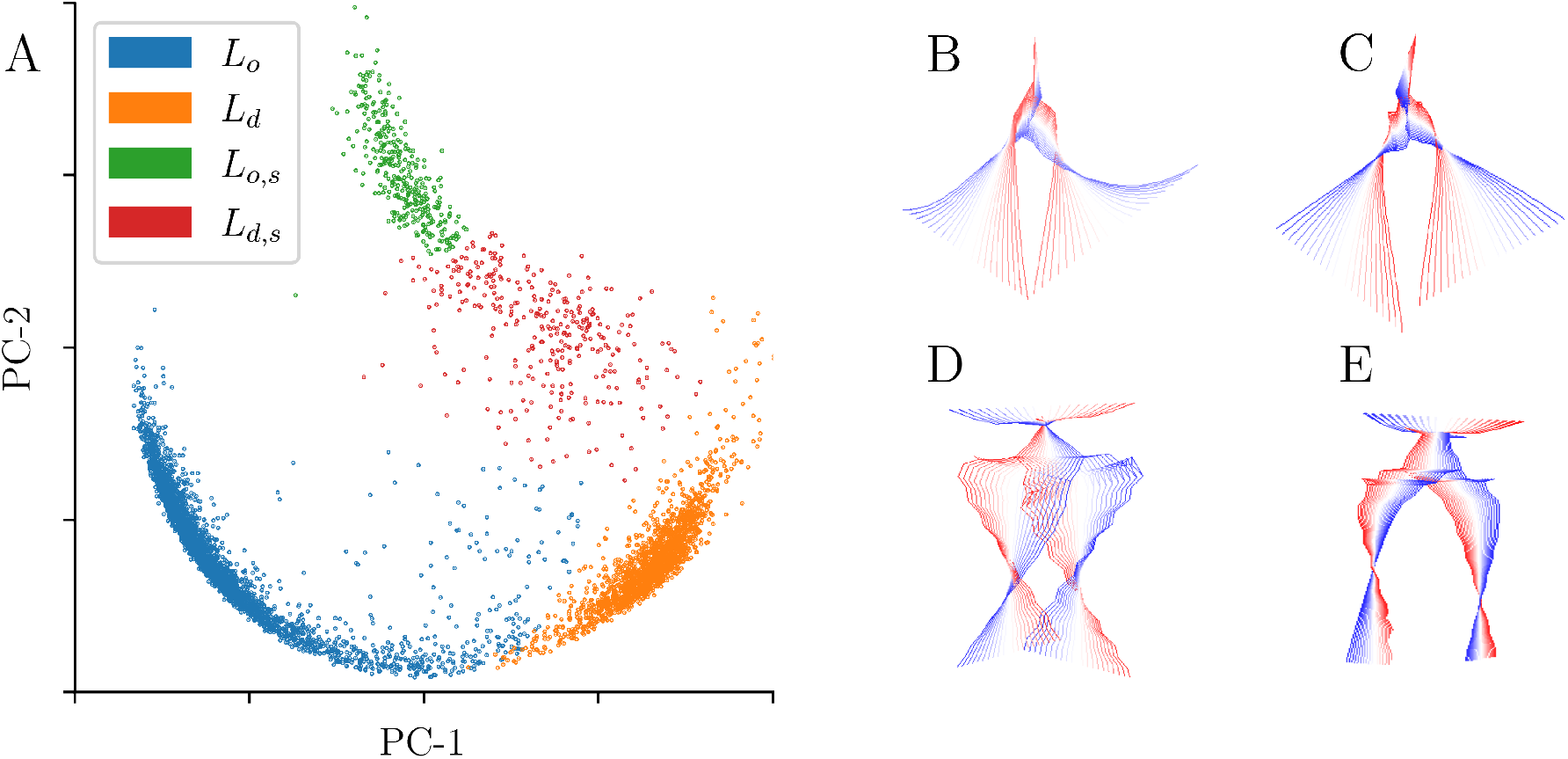
Principal component reduction of the lipid similarity matrix showing the first and second components (A). Each of the points represents a lipid and is coloured according to the cluster it was assigned to. Right: The first (most dominant) principal component from the PCA lipids analysis for *L*_d_ (B), *L*_d,s_ (C), *L*_o_ (D) and *L*_o,s_ (E). The key motion in the disordered conformations (B and C) adopt a similar scissoring/splaying tail motion. In contrast, the ordered conformations (D and E) adopt a twisting rotational motion. The colour scheme goes from red-white-blue showing the projected value of the component.

Above *T*_m_ the major component is the widely studied disordered lipid. Below *T*_m_, however, the complex, heterogeneous nature of the ripple phase becomes apparent. With the spatial ordering, laterally within a leaflet and asymmetrically across the leaflets being key to the ripple. The obvious question is: Do these four distinct conformations exist and have finite lifetimes? The number of lipids in each of the clusters as a function of temperature is shown in Figure 7A and the time dependence of the splay angle of each of the four species for one of the lipids at 315 K is show in Figure 7B. As the figure shows, all of the components are present in the ripple phase and disappear in the disordered phase. Figure 7B further shows that the components are clearly distinct. The disordered lipid (orange), switches its angle as can be expected. The other three components are very clearly present for extended times.

**Figure (7).**
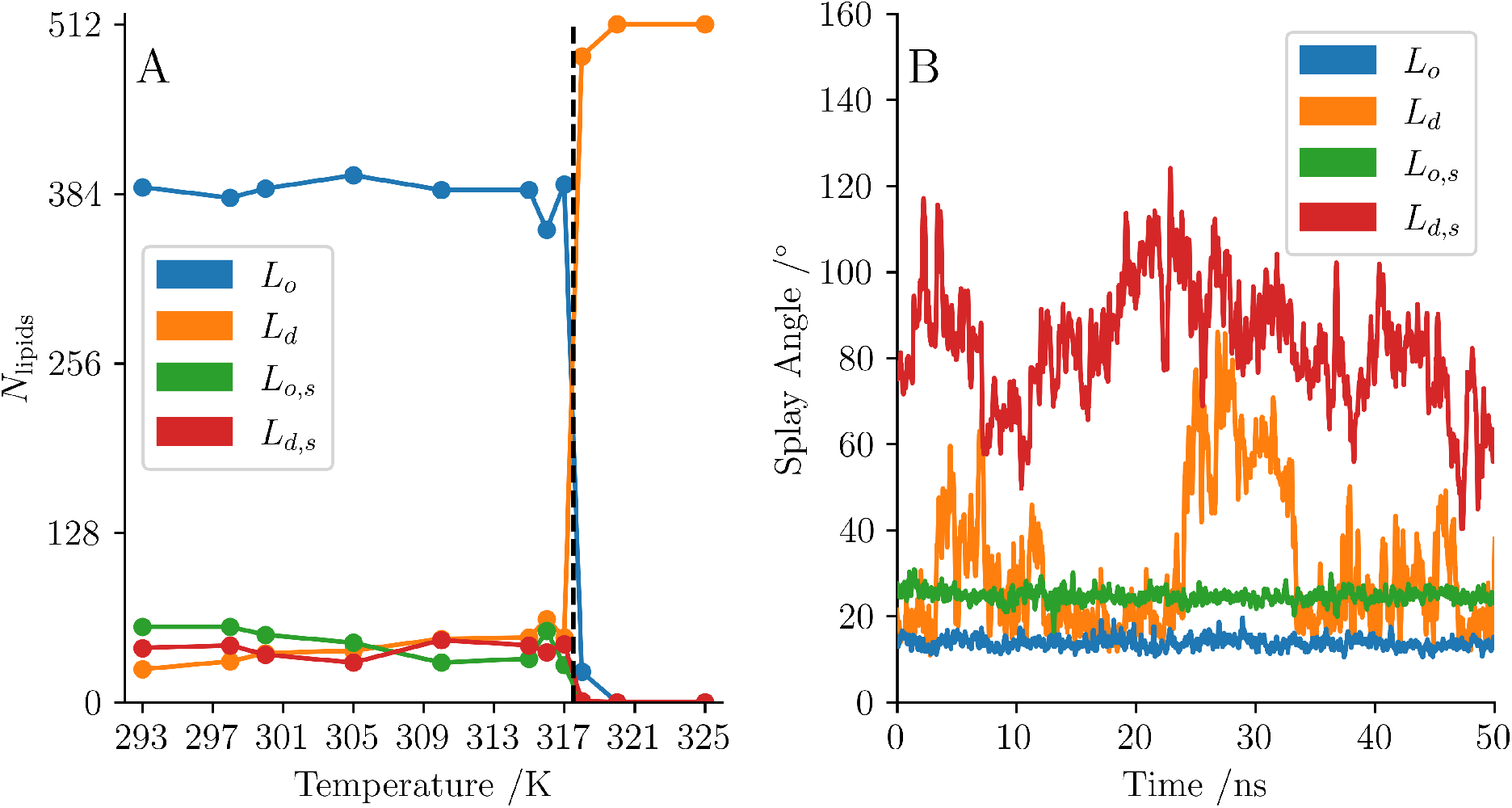
Number of lipids in each cluster at each simulation temperature (A) and the splay angle of an example lipid from each of the clusters taken from the 315K simulation (B). The black dashed line shows the phase transition temperature between 317 K and 318 K. The bulk lipid in each case is the linear lipid, ordered below the transition temperature and disordered above. Below *T*_m_ there exists disordered lipids and splayed lipids. By 320 K the only lipid component is the lipid disordered. The example splay plots show one of the key differences between the clusters, the *L*_o_ and *L*_o,s_ lipids both have a low variance whereas the *L*_d_ and *L*_d,s_ lipids are highly mobile. The *L*_d_ lipid can bee seen to transition to a splayed conformation briefly at 25 ns but returns to the more linear nature within 10 ns. Since *g*_3_ is a time averaged property the clustering is assigned based on majority of time rather than instantaneous value.

## Discussion and conclusions

In this study, we have applied a new radial-angular three-particle correlation function (*g*_3_) in combination with unsupervised machine learning to examine the phase transitions in DPPC lipid bilayers using data from atomistic molecular dynamics (MD) simulations. The analysis revealed the presence of four distinct conformational populations of lipids in the ripple phase of DPPC lipid bilayers. The expected ordered and disordered lipids are found along with their splayed equivalents. The analysis using *g*_3_ showed distinct differences in their gauche distribution and tail packing. Principal component analysis (PCA) further confirmed the existence of these four lipid groups. Finite size effects influenced smaller (128 lipid) bilayers and only three clusters were identified, with no differentiation between the ordered-splayed and disordered-splayed. The splayed lipids exist to stabilise the interface mismatch. In order to cover the interface region in smaller bilayer systems, a higher ratio of the lipids must be melted to splayed disordered. This was not the case in the small 128 lipid systems and so those systems were influenced by finite size effects.

In a very recent study Walter et al.^6^ also applied machine learning to study the main phase transition in DPPC bilayers. One fundamental difference between Walter et al. and our study is that they used supervised machine learning and trained their algorithm to distinguish between two pre-defined lipid conformations, fluid or gel. In our case, we use unsupervised machine learning and the number of lipid conformational categories was not fixed beforehand but emerged from data.

The structure of the ripple phase has remained an open question and in their 2015 study Akabori and Nagle^5^ state that the ripple structure that emerges from their scattering data does not agree with the ripple consisting of interdigitating ordered lipids as has been reported by several previous simulations, e.g. Refs.^10,11^ Akabori and Nagle dismissed the possibility of interdigitation of ordered lipids being the dominant behaviour in the thinner minor arm since the observed electron density is inconsistent with that scenario in large enough of scale. Instead, they proposed that the major arm consists of the usual ordered gel-like lipids and then hypothesized that the minor arm has five other lipid types of variable lengths and packing. Those lipids were classified as “more fluid than the gel phase” or intermediate, and suggested to exist on both sides of the minor arm. They did not elaborate these five additional lipid types further.

Our results resolve the above controversy and provide a detailed picture of the ripple structure. First of all, as discussed above, conformational clustering using unsupervised machine learning (Figure 2) revealed four distinct classes of lipids whose existence were subsequently verified by PCA. As Figures 4B,C show, disordered (orange) and disordered splayed (red) lipids cluster in the groove region of the ripple only. This is unlike to what has been suggested by Akabori and Nagle^5^ based on x-ray scattering and Khakbaz and Klauda^12^ based on their MD simulations using 72 lipids that the two sides of the ripple are symmetric in terms of lipid conformations (the two arms are asymmetric). As evident in Figure 4, the convex part has the usual ordered (blue) and some ordered splayed lipids (green) with some of them also distributed in the planar, gel-like regions. Furthermore, outside the disordered region, the minor arm consists of the usual ordered lipids (blue). Figures 4D-F also reveal that splayed lipids, both ordered and disordered splayed tend to be present only on the minor arm side of ripple. The splayed lipids act as intermediates between the ordered and disordered lipids. The disordered-splayed lipids often adopt a conformation with one of the tails relatively ordered and the other much more fluid, seemingly half transitioned, forming an interfacial layer in the ripple. As Figure 4C also shows, there is some interdigitation but that presence of the fluid phase is the key here and clarifies the picture of Akabori and Nagle.^5^ In addition, Figures 4E-F show that the idea of dislocation lines consisting of lipids in the fluid phase put forward by Heimburg^30^ over 20 years ago is indeed a very good description of the ripple phase.

The existence of the four distinct classes of lipids and their unique distribution in the convex and concave side of the ripple also shows the shortcomings of current leading theories. ^27,28,31^ While several of them do predict the ripple phase and the existence of the two arms of different lengths, none of them addresses the ripple structure or the asymmetry across the ripple. Coupling between the two leaflets and incorporation of the behaviour of individual lipids appear to be the major issues that need to be addressed in future theories.

## Methods

### Simulation methodology

Atomic-scale MD simulations of single-component DPPC lipid bilayers were performed. The systems consisted of 512 DPPC lipids and ∼20,000 TIP3P water molecules.^32^ The Gromacs software package was used in all simulations.^33^ The initial membrane structures were generated using the CHARMM-GUI Membrane Builder.^34^ The systems were first energy minimized and pre-equilibrated for 20 ns in the NPT ensemble (P=1 bar and T=325 K). The production runs were done using the V-rescale thermostat^35^ and the Parrinello-Rahman baraostat;^36^ pressure was controlled semiisotropically, 2 fs time step was used and periodic boundary conditions were applied in all directions. For the Lennard-Jones interactions, a switching function over 1-1.2 nm was used, while the particle-mesh Ewald method^37^ with a real-space cutoff of 1.2 nm was employed for electrostatics. The production simulations were ran at 293 K, 298 K, 300 K, 305 K, 310 K, 315 K, 316 K, 317 K, 318 K, 320 K and 325 K. The temperature range was chosen such that it spanned both the fluid and gel phases. A bilayer system at each temperature was simulated for 500 ns.

Robustness of the results was verified by repeating several of the simulations below *T*_m_ using different initial conditions and additionally, size dependence was assessed by simulating systems with 128 and 4,096 lipids and the usual metrics such as thickness and the area per lipid were verfied to be in agreement with previous results. The 128 lipids systems were run for 1 *µ*s each at 8 temperatures (295 K, 300 K, 305 K, 310 K, 314 K, 317 K, 318 K, and 320 K). The 4,096 lipid systems were run for 500 ns each at 3 temperatures (313 K, 315 K, 317 K). The total simulation time was 16 *µ*s.

VMD^38^ was used for visualizations, and data analysis was preformed using custom python codes, MDAnalysis,^39^ scikit-learn^40^ and scikit-image^41^ libraries. The principal components analysis (PCA) of the lipid motion was calculated using PCAlipids.^29^

## Data availability

Trajectory data is available from the authors upon request.

## Acknowledgements

The authors thank Martin Müser and Sergey Sukhomlinov for fruitful discussions regarding the application of *g*_3_ and critical reading of the manuscript. MK thanks the Natural Sciences and Engineering Research Council of Canada (NSERC) and the Canada Research Program for financial support. SharcNet and Compute Canada provided computational resources. ADRF acknowledges a postdoctoral fellowship by CONACYT-Mexico (No. 770692).

## References

(1) Tardieu, A.; Luzzati, V.; Reman, F. C. Structure and polymorphism of the hydrocarbon chains of lipids: a study of lecithin-water phases. J. Mol. Biol. 1973, 75, 711–733.

(2) Chen, S. C.; Sturtevant, J. M.; Gaffney, B. J. Scanning calorimetric evidence for a third phase transition in phosphatidylcholine bilayers. Proc. Natl. Acad. Sci. U. S. A. 1980, 77, 5060–5063.

(3) Sun, W. J.; Tristram-Nagle, S.; Suter, R. M.; Nagle, J. F. Structure of the ripple phase in lecithin bilayers. Proc. Natl. Acad. Sci. U. S. A. 1996, 93, 7008–7012.

(4) Sengupta, K.; Raghunathan, V. A.; Katsaras, J. Novel structural features of the ripple phase of phospholipids. Europhys. Lett. 2000, 49, 722.

(5) Akabori, K.; Nagle, J. F. Structure of the DMPC lipid bilayer ripple phase. Soft Matter 2015, 11, 918–926.

(6) Walter, V.; Ruscher, C.; Benzerara, O.; Marques, C. M.; Thalmann, F. A machine learning study of the two states model for lipid bilayer phase transitions. Phys. Chem. Chem. Phys. 2020, 22, 19147–19154.

(7) Mouritsen, O. G. Studies on the lack of cooperativity in the melting of lipid bilayers. Biochim. Biophys. Acta 1983, 731, 217–221.

(8) Lemmich, J.; Mortensen, K.; Ipsen, J. H.; Honger, T.; Bauer, R.; Mouritsen, O. G. Pseudocritical behavior and unbinding of phospholipid bilayers. Phys. Rev. Lett. 1995, 75, 3958–3961.

(9) Kuklin, A.; Zabelskii, D.; Gordeliy, I.; Teixeira, J.; Brûlet, A.; Chupin, V.; Cherezov, V.; Gordeliy, V. On the Origin of the Anomalous Behavior of Lipid Membrane Properties in the Vicinity of the Chain-Melting Phase Transition. Sci. Rep. 2020, 10, 5749.

(10) de Vries, A. H.; Yefimov, S.; Mark, A. E.; Marrink, S. J. Molecular structure of the lecithin ripple phase. Proc. Natl. Acad. Sci. U. S. A. 2005, 102, 5392–5396.

(11) Lenz, O.; Schmid, F. Structure of Symmetric and Asymmetric “Ripple” Phases in Lipid Bilayers. Phys. Rev. Lett. 2007, 98, 058104.

(12) Khakbaz, P.; Klauda, J. B. Investigation of phase transitions of saturated phosphocholine lipid bilayers via molecular dynamics simulations. Biochim. Biophys. Acta (BBA) - Biomembranes 2018, 1860, 1489–1501.

(13) Sukhomlinov, S. V.; Müser, M. H. A mixed radial, angular, three-body distribution function as a tool for local structure characterization: Application to single-component structures. J. Chem. Phys. 2020, 152, 194502.

(14) Sukhomlinov, S. V.; Müser, M. H. Stress Anisotropy Severely Affects Zinc Phosphate Network Formation. Tribol. Lett. 2021, 69, 89.

(15) Smith, S. O.; Hamilton, J.; Salmon, A.; Bormann, B. J. Rotational Resonance NMR Determination of Intra-and Intermolecular Distance Constraints in Dipalmitoylphosphatidylcholine Bilayers. Biochemistry 1994, 33, 6327–6333.

(16) Wang, Z.; Bovik, A. C.; Sheikh, H. R.; Simoncelli, E. P. Image quality assessment: from error visibility to structural similarity. IEEE Trans. Image Process. 2004, 13, 600–612.

(17) Ziolek, R. M.; Smith, P.; Pink, D. L.; Dreiss, C. A.; Lorenz, C. D. Unsupervised Learning Unravels the Structure of Four-Arm and Linear Block Copolymer Micelles. Macromolecules 2021, 54, 3755–3768.

(18) Smith, P.; Quinn, P. J.; Lorenz, C. D. Two Coexisting Membrane Structures Are Defined by Lateral and Transbilayer Interactions between Sphingomyelin and Cholesterol. Langmuir 2020, 36, 9786–9799.

(19) van der Maaten, L.; Hinton, G. Visualizing Data using t-SNE. J. Mach. Learn. Res. 2008, 9, 2579–2605.

(20) Schubert, E.; Sander, J.; Ester, M.; Kriegel, H. P.; Xu, X. DBSCAN Revisited, Revisited: Why and How You Should (Still) Use DBSCAN. ACM Trans. Database Syst. 2017, 42, 1–21.

(21) Kriegel, H.-P.; Schubert, E.; Zimek, A. The (black) art of runtime evaluation: Are we comparing algorithms or implementationsã Knowl. Inf. Syst. 2017, 52, 341–378.

(22) Biltonen, R. L.; Lichtenberg, D. The use of differential scanning calorimetry as a tool to characterize liposome preparations. Chem. Phys. Lipids 1993, 64, 129–142.

(23) Ivanova, V. P.; Heimburg, T. Histogram method to obtain heat capacities in lipid monolayers, curved bilayers, and membranes containing peptides. Phys. Rev. E 2001, 63, 041914.

(24) Chen, W.; Duša, F.; Witos, J.; Ruokonen, S.-K.; Wiedmer, S. K. Determination of the Main Phase Transition Temperature of Phospholipids by Nanoplasmonic Sensing. Sci. Rep. 2018, 8, 1–11.

(25) Sun, L.; Böckmann, R. A. Membrane phase transition during heating and cooling: molecular insight into reversible melting. Eur. Biophys. J. 2018, 47, 151–164.

(26) Walter, V.; Ruscher, C.; Gola, A.; Marques, C. M.; Benzerara, O.; Thalmann, F. Ripple-like instability in the simulated gel phase of finite size phosphocholine bilayers. Biochim. Biophys. Acta – Biomembranes 2021, 183714.

(27) Lubensky, T. C.; MacKintosh, F. C. Theory of “ripple” phases of lipid bilayers. Phys. Rev. Lett. 1993, 71, 1565–1568.

(28) Kamal, M. A.; Pal, A.; Raghunathan, V. A.; Rao, M. Theory of the asymmetric ripple phase in achiral lipid membranes. Europhys. Lett. 2011, 95, 48004.

(29) Buslaev, P.; Gordeliy, V.; Grudinin, S.; Gushchin, I. Principal Component Analysis of Lipid Molecule Conformational Changes in Molecular Dynamics Simulations. J. Chem. Theory Comput. 2016, 12, 1019–1028.

(30) Heimburg, T. A Model for the Lipid Pretransition: Coupling of Ripple Formation with the Chain-Melting Transition. Biophys. J. 2000, 78, 1154–1165.

(31) Goldstein, R. E.; Leibler, S. Model for lamellar phases of interacting lipid membranes. Phys. Rev. Lett. 1988, 61, 2213–2216.

(32) MacKerell, A. D. et al. All-atom empirical potential for molecular modeling and dynamics studies of proteins. J. Phys. Chem. B 1998, 102, 3586–3616.

(33) Abraham, M. J.; Murtola, T.; Schulz, R.; Páll, S.; Smith, J. C.; Hess, B.; Lindahl, E. GROMACS: High performance molecular simulations through multi-level parallelism from laptops to supercomputers. SoftwareX 2015, 1-2, 19–25.

(34) Lee, J. et al. CHARMM-GUI Input Generator for NAMD, GROMACS, AMBER, OpenMM, and CHARMM/OpenMM Simulations Using the CHARMM36 Additive Force Field. J. Chem. Theory Comput. 2016, 12, 405–413.

(35) Bussi, G.; Donadio, D.; Parrinello, M. Canonical sampling through velocity rescaling. J. Chem. Phys. 2007, 126, 014101.

(36) Parrinello, M.; Rahman, A. Polymorphic transitions in single crystals: A new molecular dynamics method. J. Appl. Phys. 1981, 52, 7182–7190.

(37) Essman, U.; Perera, L.; Berkowitz, M. L.; Darden, T.; Lee, H.; Pedersen, L. G. A smooth particle mesh Ewald method. J. Chem. Phys. 1995, 103, 8577–8592.

(38) Humphrey, W.; Dalke, A.; Schulten, K. VMD: Visual molecular dynamics. J. Mol. Graph. 1996, 14, 33–8, 27–8.

(39) Michaud-Agrawal, N.; Denning, E. J.; Woolf, T. B.; Beckstein, O. MDAnalysis: A toolkit for the analysis of molecular dynamics simulations. J. Comput. Chem. 2011, 32, 2319–2327.

(40) Pedregosa, F. et al. Scikit-learn: Machine Learning in Python. J. Mach. Learn. Res. 2011, 12, 2825–2830.

(41) van der Walt, S.; Schönberger, J. L.; Nunez-Iglesias, J.; Boulogne, F.; Warner, J. D.; Yager, N.; Gouillart, E.; Yu, T.; the scikit-image contributors, scikit-image: image processing in Python. PeerJ 2014, 2, e453.

